# Data-theoretical Synthesis of the Early Developmental Process

**DOI:** 10.1101/282004

**Authors:** Bradly Alicea, Richard Gordon, Thomas E. Portegys

## Abstract

Biological development is often described as a dynamic, emergent process. This is evident across a variety of phenomena, from the temporal organization of cell types in the embryo to compounding trends that affect large-scale differentiation. To better understand this, we propose combining quantitative investigations of biological development with theory-building techniques. This provides an alternative to the gene-centric view of development: namely, the view that developmental genes and their expression determine the complexity of the developmental phenotype. Using the model system *Caenorhabditis elegans*, we examine time-dependent properties of the embryonic phenotype and utilize the unique life-history properties to demonstrate how these emergent properties can be linked together by data analysis and theory-building. We also focus on embryogenetic differentiation processes, and how terminally-differentiated cells contribute to structure and function of the adult phenotype. Examining embryogenetic dynamics from 200 to 400 minutes post-fertilization provides basic quantitative information on developmental tempo and process. To summarize, theory construction techniques are summarized and proposed as a way to rigorously interpret our data. Our proposed approach to a formal data representation that can provide critical links across life-history, anatomy and function.

## Introduction

The understanding of development as a dynamic, emergent process stands at odds with our current understanding of large-scale developmental patterns. While there have been attempts to characterize these patterns using physical laws [1, 2], accessing these phenomena with formal logical descriptions is also useful for purposes of both modeling and connections to molecular mechanisms. In this paper, we will build towards bridging patterns observed as cells differentiate in an embryo with the potential to construct theories and mathematical applications. This will bring us closer to understanding how to construct theories of developmental biology, particularly theories that explain and perhaps predict the emergence of developmental phenotypes.

Understanding embryogenetic systems in this way is not attainable using a gene-centric approach. To overcome this limitation of more traditional developmental biology research, we have engaged in work that demonstrates some of these relational processes during embryogenesis. This investigation will focus on a specific question: in what order do distinct cells emerge within and between tissue types at multiple time points in pre-hatch morphogenesis? Ultimately, such explorations of data will be useful in building compositional models of development where we consider how various component elements are combined to build a whole embryo. Additionally, we wish to provide a means to building a systems theory of development, one that will lead us to more formal representations of the data. We propose that developmental processes, particularly in deterministic model organisms such as *C. elegans*. Such a system is useful for casting our analysis of embryonic development as a set of relationships amenable to theory-building techniques.

This paper will proceed by introducing the reader to cellular-level alternatives to reductionism in the study of development, an analysis of differentiation into terminal adult cell types, and an analysis of early development. We assume that a temporal analysis of early stages in the differentiation process (in this case, 200 to 400 minutes of *C. elegans* embryogenesis) can reveal much about the emergence of larger-scale processes and structures in the developmental phenotype [3, 4]. The first and second points provide a means to better understand the connection between developmental cell lineages and their differentiated descendant cells. A discussion and synthesis of this analysis is presented, followed by the potential use of theory-building to interpret these results. While this paper features little in terms of formal mathematics, it can nonetheless be greatly useful to mathematical biologists, particularly those interested in the nematode *Caenorhabditis elegans*. However, we provide hypothetical explanations that could be checked using mathematical models. These explanations might serve as heuristics for further research, including applications of more formal frameworks such as category theory.

### Cellular-level Alternatives to Gene-centrism

Our approach is based on quantitative characterization and phenomenological modeling of development. In this paper, both digital approaches to morphogenesis as well as cellular-level models serve as alternatives to a gene-centric approach. Through the use of cellular-level computational models, we can account for short-range interactions such as paracrine signaling and physical interactions. We can also combine the results from various primary data sets into a model of selective interactions between cells and regions of the embryo. Our work on establishing an “interactome” for *C. elegans* embryos [5] is an example of the value of such local-to-global information. There is also value in establishing frameworks for multiple types of data, which might lead to inference or insight down the road. The availability of both molecular and cellular data at the single cell level in *C. elegans* provides a unique opportunity to ask questions such as how the physiology of embryogenesis unfolds in space.

### Theory-building for Data Science

Quantitative data analysis is inseparable from theory-building. The systems alternative to gene-centrism we propose very much depends upon a theoretical interpretation of the data. Lack of theory hinders progress in two ways: we cannot understand how things relate to one another, and we cannot develop a framework for intervention [6]. To overcome these obstacles, theory development must transcend the hypothetico-deductive method. This makes the practice of modern biomedical science somewhat incompatible. Yet theory construction can encourage rigor in the standards in computational biology and secondary data analysis. A theoretical analysis should enable four properties from the data: conceptual definitions, domain limitations, relationship-building, and predictions. This theory must then allow for properties such as uniqueness, parsimony, conservation, generalizability, fecundity, internal consistency, empirical riskiness, abstraction, and similarities between different domains [7]. Corley and Gioia [8] have proposed two additional goals of theory-building. The originality criterion refers to theories allowing for revelatory insights while also supporting the inherently incremental nature of investigation.

Theory serves two purposes: discerning and anticipating what we need to know, and influencing the intellectual theory-building framing and dialogue [8]. Additionally, the utility of a theory requires it to be both practical (plausible according to the working of the natural world) and accessible to investigation via formal scientific methods. A prospective theory should involve at least three types of subjects and two types of investigation [9]. Generally speaking, theories organize a variety of objects, sources of knowledge, and kinds of knowledge to provide a reasonable characterization of the system in question. This includes synthesizing data in the service of characterizing internal structure of the system in question [10]. While this is a difficult problem that can lead to seemingly unending complexity [11], it is nevertheless necessary for proper interpretation.

As theory is not merely about organizing sets of relationships, we also need a formal investigatory framework. In understanding the relationship between objects as opposed to the nature of objects, theories can either be analytic or synthetic [9]. While analytic approaches utilize mathematical and philosophical tools, synthetic approaches involve bringing together a variety of different data sources to propose new principles for the system in question. The current study provides an *a priori* synthetic perspective [10], as we provide a theory of content, which adds knowledge to future analytical efforts.

### Analysis of Development

In this section, we will discuss current initiatives and future directions in the analysis of development. This includes a discussion of developmental cell organization, an overview of developmental cell lineage and differentiation trees, segmentation/partitioning of imaging data, and the extraction of developmental dynamics.

#### Developmental Cell Organization

*C. elegans* has a mode of development called *mosaic development*. While this is different from embryonic *regulative development* in amphibians and mammals, in which many cells appear to have equivalent roles [12], there are many other examples of mosaic development throughout the tree of life. Mosaic development is a process whereby most developmental cells have determined fates. After the initial cleavages in *C. elegans*, there are six founder cells (*AB, C, D, E, MS, P4*) which go on to produce specialized lineages of cells with no variation across individuals. These sublineages contribute to various tissue and anatomical structures in the adult worm. *C. elegans* is eutelic, which means that there is a fixed number of somatic cells in the adult.

#### Developmental Cell Lineage Tree

The *C. elegans* lineage tree [13] describes the lineal order of descent for all developmental cells from the one-cell stage to terminal differentiation or cell death. The lineage tree is ordered along the anterior-posterior axis of the worm [14], and describes the lineage of descent leading to all cells in the adult worm. Sublineages (descendants of the founder cells) consist of multiple layers of cells, which diversify at fixed times before becoming terminally differentiated cell types, also at fixed times. The timings of these division events are rather uniform across layers of the tree, although there are some notable exceptions.

#### Developmental Cell Differentiation Tree

Lineage trees have been proven to be adequate data structures for organizing information about developmental cell descent. However, different modes of data organization and analysis are possible. In nematodes, differentiation trees allow us to relate the binary, mostly asymmetric cell divisions to the broader context of embryonic tissue differentiation. Alternative methods include meta-Boolean models [15], complex networks [5], algorithmic complexity [16], and scale-invariant power laws [17]. One method that relies upon simply reorganizing the lineage tree by the occurrence of differentiation waves is called differentiation tree analysis [12].

## Data and Visualization – Methods

### Pre-Hatch Morphogenesis and Timepoints

Pre-hatch morphogenesis in *C. elegans* is the period from fertilization to 400 minutes of embryogenesis at 25°C. We begin sampling intervals at 200 minutes, at which time the only terminally differentiated cells are the germ cells. Our sampled time points are at intervals of 5 minutes from 200 to 300 minutes of embryogenesis, and at intervals of 50 minutes between 300 and 400 minutes of embryogenesis. This gives us a total of 23 time points: 21 points over the 200 to 300 minute interval, and two points post-300 minutes (350 and 400). Many of the major terminal cell types emerge between 200 and 300 minutes, making sparse sampling of the post-300 minute period adequate.

### Timed Cell Lineage Data

Timed cell lineage data were acquired courtesy of Nikhil Bhatla and his lineage tree application (http://wormweb.org/celllineage). Cells represented in an embryo at a given time are determined by first calculating the lifespan of each cell in the lineage tree (e.g. the time at which each cell is born and either divides or dies), and then identifying all cells alive at a given time. Terminally-differentiated cells were assumed never to die, unless specified by the data.

### Developmental and Terminally-Differentiated Cells

In this set of analyses, we focus on two distinct cellular states: developmental cells and terminally-differentiated cells. Developmental cells are those that generally produce daughter cells, unless they are products of a terminal cell division. In *C. elegans*, developmental cells are part of an invariant cell lineage in which fate acquisition is predetermined [13]. By contrast, terminally-differentiated cells are those which have both reached a terminal division and have acquired a terminal functional state. For example, if a cell lineage results in muscle cells, then developmental cells will divide until they stop dividing and acquire the muscle fate. The role of post-embryonic plasticity in the many facets of cell identity [18] is unknown, so one caveat of our analyses is that terminally-differentiate cells observed in the embryo or larval worm are not identical to their adult counterparts.

### Cell Functional Annotations

The annotation of each terminally-differentiated cell’s function was acquired courtesy of Stephen Larson and Mark Watts from the PyOpenWorm project (https://github.com/openworm/PyOpenWorm). Annotations were matched to each cell name using the Sulston nomenclature system [13], which resulted in a series of annotated cells (*C_i_*) at time *t*. This also resulted in a function *t_i_*(*x*) for each cell (1, 2,……, *x*). Text mining was then used to determine a given cell’s functional class. The birth of functional classes over developmental time was done using a binary classifier.

### Functional Classes and Families

To look at differences within and between groups of terminally-differentiated cells, we used a two-tiered classification scheme. This consisted of functional classes and families. Functional classes are based on annotation identities, which are extracted as keywords found in the list of annotations. Families are groups of cells with the same first letter or prefix in their nomenclature identity (e.g. all cells with the nomenclature identity *hyp* belong to the same family). In the heat maps (Supplemental Figures 1-3), these categories are shown to largely overlap.

### CAST (Cell Alignment Search Tool)

The original methodology for the CAST Alignment is shown in [19]. In this analysis, we calculate pairwise CAST alignment for the current time point and the next time point. The CAST alignment yields an alignment score, which is divided into the maximum possible score to yield the CAST coefficient. The maximum possible score is equivalent to the length of the cell list for the next time point (the longer cell list of the two cell lists in the pairwise comparison). This value of this coefficient can range from −1 to 1, and allows for a time-series of these pairwise comparisons to be compared.

### Cluster Analysis and Information Content

A hierarchical cluster analysis was conducted using R version 3.3.1. The data were visualized using Rstudio 0.99. The cluster vector matrix was extracted, transposed, and vectorized using SciLab 5.5.2. The cluster vector is then used to determine how many cells from each family (n=26) belong to each cluster. This allows for the Shannon Information for each cell family to be calculated.

### Data Accessibility

All processed data is available in the Supplemental Files, which are also archived at the Open Science Framework (https://osf.io/u8abh/). Data for the DevoWorm lineage tree (https://github.com/balicea/DevoWorm/tree/master/Lineage%20Tree%20DB) and differentiation tree (https://github.com/balicea/DevoWorm/tree/master/Differentiation%20Tree%20Dataset) data sets are archived on Github.

Demos are also available in the form of interactive Jupyter Notebooks. These notebooks demonstrate concepts such as time-series for differentiation processes (https://github.com/devoworm/devoworm.github.io/blob/master/Differentiation-Time-series-in-C-elegans-and-Drosophila.ipynb), the process of representing differentiation trees as binary graphs (https://github.com/devoworm/devoworm.github.io/blob/master/Differentiation%2BTree%2Bas%2BGraphs.ipynb), and the concept of Embryo Space (https://github.com/devoworm/devoworm.github.io/blob/master/Embryo-Space-Concept-in-C-elegans.ipynb).

## Data and Visualization – Results

We conducted an analysis of publicly available data demonstrating the unfolding of adult morphology during embryogenesis. The first step in the analysis is to show the number of developmental and terminally-differentiated cells from 200-400 minutes. These data are available in tabular form for annotated nomenclature identities (Supplemental File 1) and for five distinct somatic cell types (Supplemental File 2). A more finely sampled demographic representation of the 200-300 minute interval shown in Figure 1 and Supplemental Figure 1. Perhaps more surprisingly is that developmental cells are added along with an increasing number of terminal-differentiation cells until around 250 minutes of embryogenesis. At around the same time, there is an inflection point for developmental cell number and an increase in the number of terminally-differentiated cells in the embryo.

**FIGURE 1.**
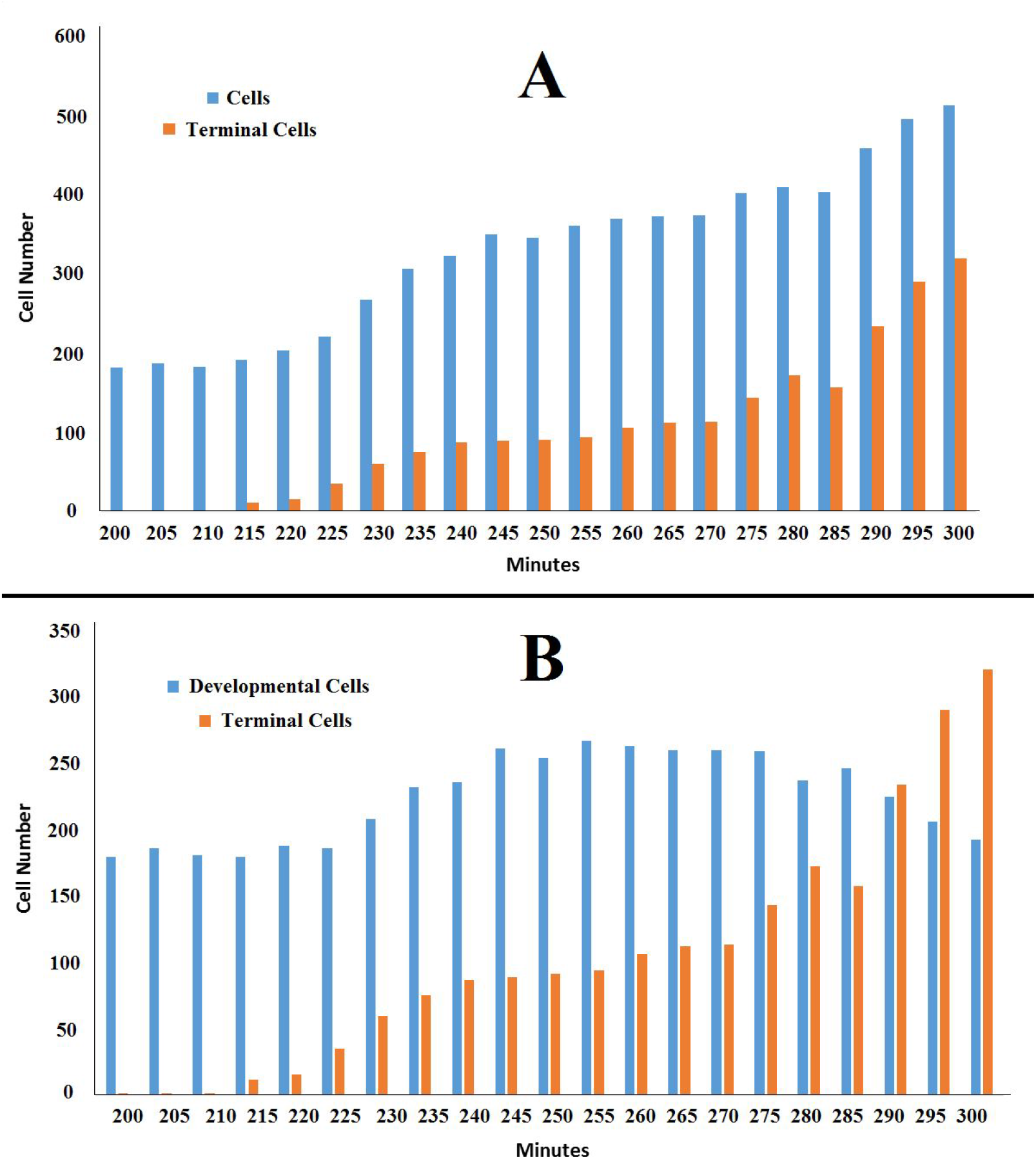
The ratio of all cells to terminally-differentiated cells (A, top) and developmental cells to terminally-differentiated cells (B, bottom) at 5 minute intervals from 200 to 300 minutes of embryogenesis. Data collected from embryos raised at 25°C.

In general, Figure 1 also provides two critical pieces of information about developmental dynamics. Figure 1A shows that the number of cells increases 2.5-fold over that 100 minute interval. One consequence of this finding suggests a periodicity in the rate of expansion in the number of cells of the embryo. In Figure 1A, it appears that there are periods of relative stasis and periods where the rate of division and differentiation increase. One of these apparent periods of stasis is from 235 to 270 minutes for terminally-differentiated cells, and 245 to 270 minutes for all cells. This includes both developmental and terminally-differentiated cells, so the difference in stasis time is likely due to changes in developmental cell number.

Figure 1 also demonstrates how the number of terminally-differentiated cells exceeds the number of developmental cells in the period from 285 to 290 minutes (Figure 1B). After 285 minutes, the *C. elegans* embryo is increasingly dominated by terminally-differentiated cells, as the number of developmental cells decreases. There are roughly the same number of developmental cells at the beginning and end of this time interval. However, in the middle of this interval (from roughly 230 to 285 minutes), there is an increase in the number of developmental cells. This is probably to feed the large increase in terminally-differentiated cells in the subsequent time periods (from roughly 285 to 350 minutes).

Both Figure 2 and Supplemental Figure 2 demonstrate these changes in the number of developmental cells, but broken out by sublineage. In Supplemental Figure 2A, we demonstrate both the number of cells in AB and MS, and the transient increase in developmental cells for sublineages AB and MS. Supplemental Figure 2B shows the number of cells in the C, D, and E sublineages. Interestingly, the fluctuation pattern demonstrated in Supplemental Figure 2A does not occur in sublineage C, and is hard to identify in sublineages D and E (Supplemental Figure 2B).

**FIGURE 2.**
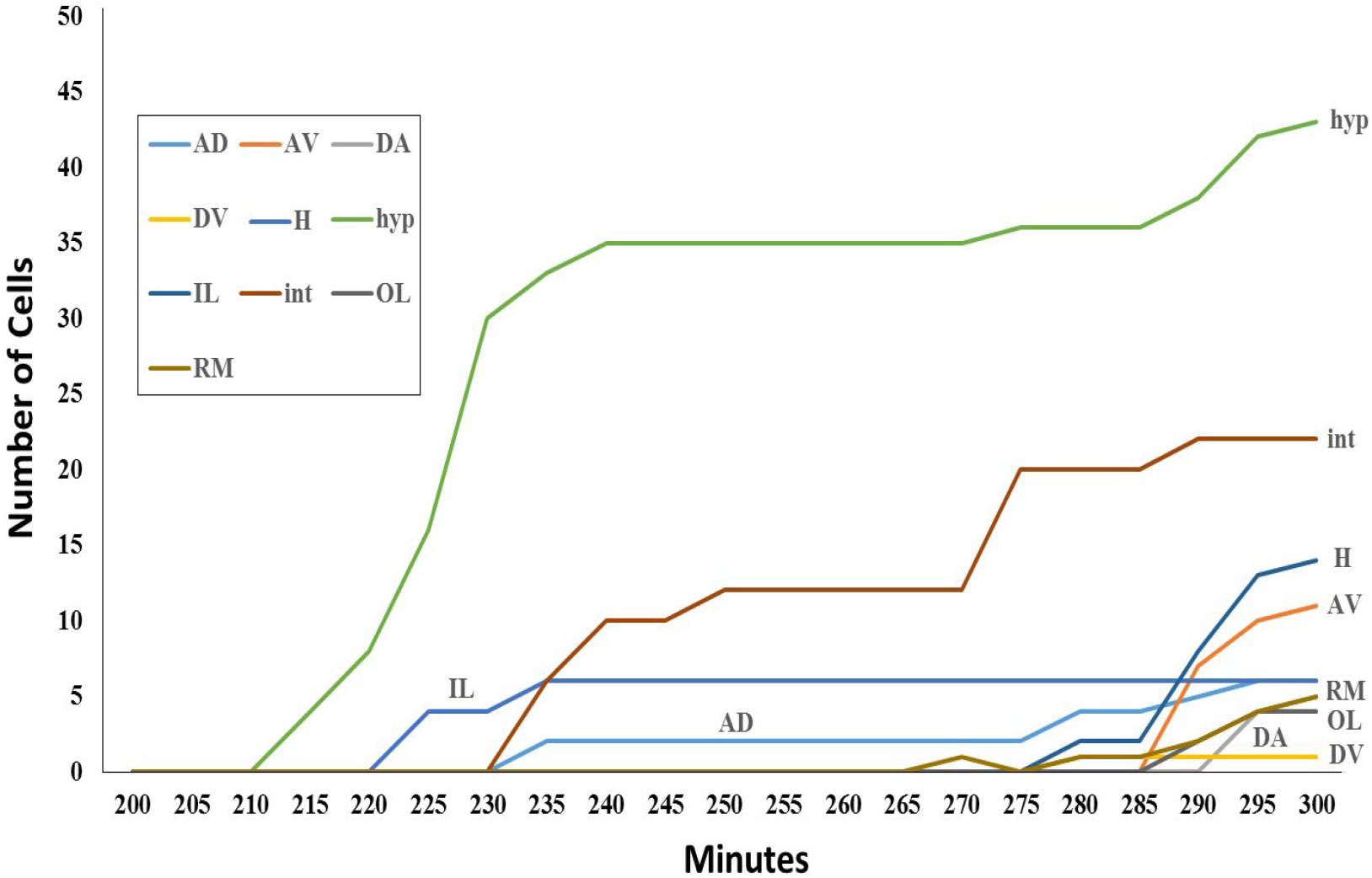
Number of cells for selected families of terminally-differentiated cells for 5-minute intervals over 200-300 minutes of embryogenesis. *AD* = Amphid cell, anterior deirid; *AV* = *Neurons*, interneurons; *DA* = Ventral motorneuron; *DV* = Ring interneurons; *H* = Seam hypodermal cell; *hyp* = Hypodermal cells; *IL* = Inner labial; *int* = Intestinal cell; *OL* = Outer labial; *RM* = Ring motorneuron/interneuron. Data collected from embryos raised at 25 °C.

Now we turn to changes in the number of terminally-differentiated cells over time, particularly as broken down by specific cell families (e.g. nomenclature identities sharing the same prefix). In Figure 2, we can see that increases in the number of cells in each family differ in both rate of increase and time of origin. Hypodermal (*hyp*) cells begin to terminally-differentiate first, followed by amphid (*AD*), inner labial (*IL*), and intestinal (*int*) cells. Up to 300 minutes, the majority of cells of the subsample in Figure 6 are hypodermal and intestinal cells. Using 200 minutes as a baseline for the earliest possible terminal differentiation, *hyp*, *AD*, *IL*, and *int* cells are what we consider to be early emerging cells.

While different terminally-differentiated cell families emerge at different times, a more relevant question with respect to organ and tissue formation is how do different co-functional cell types compare in terms of their rate and time of differentiation. The heat map in Supplemental Figure 3 contains all terminally-differentiated cells present up to 400 minutes of embryogenesis. Each color represents a specific nomenclature identity which corresponds to a specific functional class (e.g. neuron, hypodermal, muscle) of an individual cell. These relationships are explored for three different pairs of cell types in Figures 3–5. Figures 3 (Hypodermal and Interneuronal), 4 (Neuronal and Syncytium), and 5 (Muscle and Synticium) show subsets of the main heat map contiguously, which compares two classes of terminally-differentiated cell in a continuous fashion.

**FIGURE 3.**
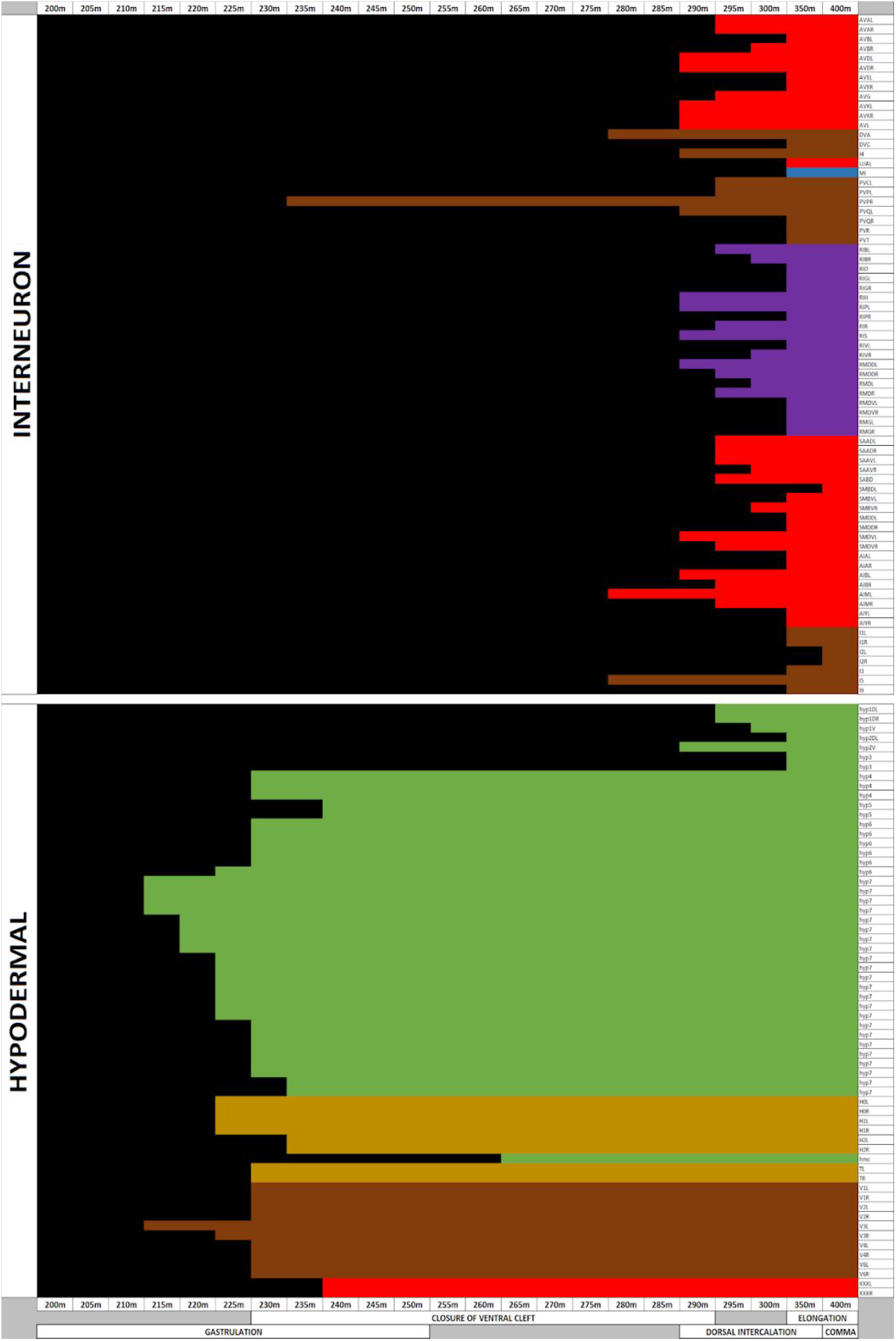
Colored chart showing the emergence of terminally differentiated cells in *C. elegans* from 200 to 400 minutes of embryogenesis showing only a comparison of hypodermal and interneuronal cells. Data collected from embryos raised at 25°C.

The heat map visualization gives us a rough guide to the amount of heterogeneity in each functional class with respect to time of birth. For some functional classes (nomenclature identity “*h*”), the birth of cells overwhelmingly occurs early in the 200 to 400 minute window of development. In other functional classes (nomenclature identity “*i*”), there is structured variation with respect to birth time. A third set of functional classes (nomenclature identities “*A*” and “*M*”) also demonstrate variation in timing between cells. Supplemental File 3 shows the descriptive statistics for each family and functional class of cell present in the embryo up to 400 minutes of embryogenesis.

As they both represent cell types that form the emerging connectome, a comparison of neurons and interneurons in terms of their emergence time is warranted. In Supplemental Figure 4, we compare the joint distribution of emergence time for three types of differentiated cells (neurons, interneurons, and hypodermal) in two comparisons. In Supplemental Figure 4A, we directly compare neurons and interneurons. Supplemental Figure 4B shows an evaluation of interneurons and hypodermal cells in the same manner. In the case of Supplemental Figure 4A, neurons merge in a bimodal fashion (with a majority of terminally-differentiated neurons being born from 290-400 minutes). By contrast, interneurons seem to almost always emerge after 280 minutes. Critically, there is an overlap in terms of terminal-differentiation between the two cell types. This may reveal an interdependency between the two cell types. By contrast, Supplemental Figure 4B shows a difference in mode between interneurons and hypodermal cells, with their frequency of emergence being almost inverse with respect to the 200 to 400 minute time interval.

The “Interneuron” functional class in Figure 4 shows the phenomenon of structured variation in more detail. In the heat map, the emergence of cells at different points in time look like jagged teeth across the cell identity (vertical) axis. This represents the birth of axial variants of the same cell type at slightly different points in time.

**FIGURE 4.**
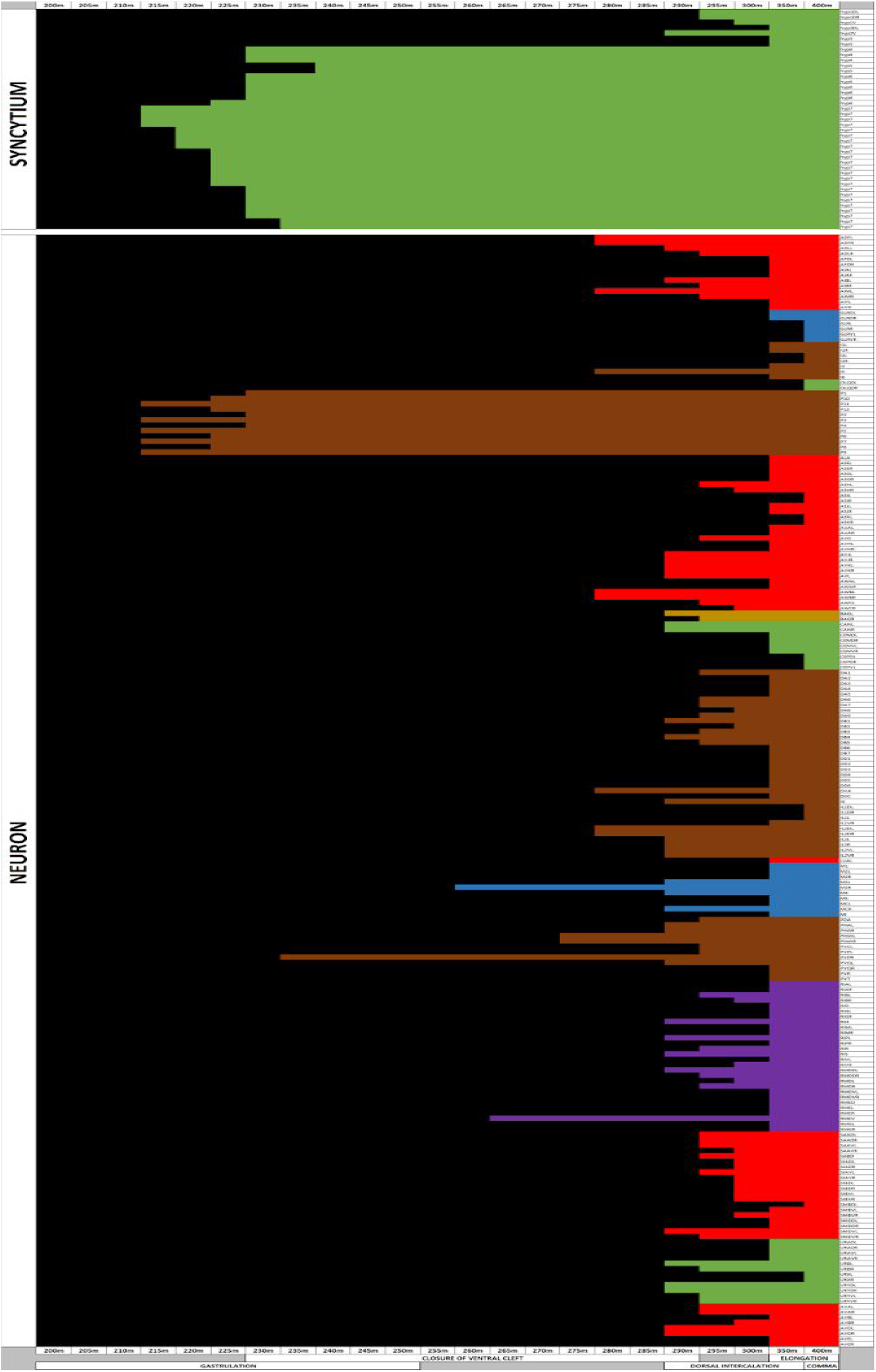
Colored chart showing the emergence of terminally differentiated cells in *C. elegans* from 200 to 400 minutes of embryogenesis showing only a comparison of neuronal and synticial cells. Data collected from embryos raised at 25°C.

Figure 5 shows the relationship between syncytium and muscle cells. For the most part, syncytium emerges earlier in time than muscle cells. However, there is a group of embryonic body wall (*mu bod*) cells born just after the first wave of syncytia. More closely resembling the timing of neuronal cells, these syncytia differentiate much earlier than the other embryonic body wall cells in our data set.

**FIGURE 5.**
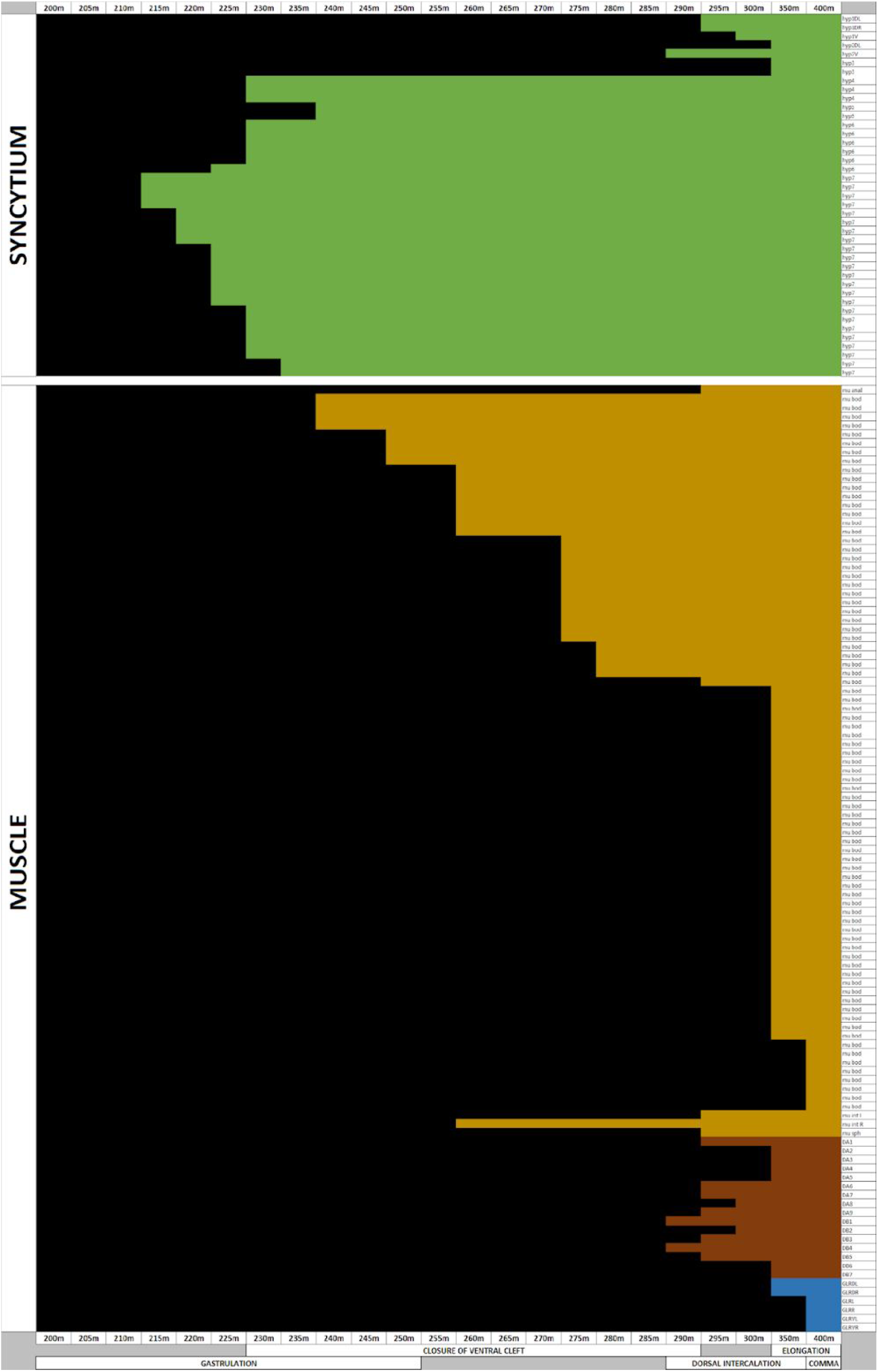
Colored chart showing the emergence of terminally differentiated cells in *C. elegans* from 200 to 400 minutes of embryogenesis showing only a comparison of muscle and synticial cells. Data collected from embryos raised at 25°C.

Looking more closely at axial variants with the same identity, we can see that while some axial variants emerge at the same time (e.g. *AIAL* and *AIAR*, right/left homologues of amphid interneurons), others emerge 5-15 minutes apart. Examples of these include *SMBDL* and *SMBVL* (dorsal/ventral homologues of ring/ motor interneurons) and *RIPR* and RIPL (right/left homologues of ring/pharynx interneurons).

We can also look at the relationship between the time of birth and number of cells per functional class. To discover patterns in these data, we conducted a hierarchical cluster analysis on the birth times for each terminally-differentiated cell. Supplemental File 4 provides an overview of the relationship between cluster membership and nomenclature family. This provides us with a set of 17 distinct clusters which we can use to classify each cell. Given this information, we asked whether cells from the same nomenclature family belonged to the same cluster. Supplemental Figure 5 shows the variation in information content across nomenclature families. The closer the value is to 1.0, the greater the information (e.g. cells from a single family are represented in a greater number of clusters).

Supplemental Figure 5 demonstrates that there are four types of nomenclature families: 1) relatively high information content with few members, 2) relatively low information content with few members, 3) relatively high information content with many members, and 4) relatively low information content with many members. This can be determined quantitatively by classifying the families based on whether their information content and cell number is above or below the median value of each. Using this method, we can determine the number of families in each category and their exemplars. Exemplars of Type 1 (1 family out of 26) include family U. Exemplars of Type 2 (12 families out of 26) include families B, E, G, and rect. Exemplars of Type 3 (11 families out of 26) include families D, M, mu, and V. Exemplars of Type 4 (1 family out of 26) include family C.

Finally, we can examine the series of terminally-differentiated cells that emerge at different time points as a CAST alignment [20]. CAST alignments provide an assessment of gaps in series of functionally-related cells as well as potential periods of stasis in the differentiation process (Supplemental File 5). Supplemental Figure 6 shows us the pattern for the 200 to 400 minutes of *C. elegans* embryogenesis time-series. In this time-series, we see a large fluctuation in the CAST coefficient between the 205-210 minute interval and the 240-245 minute interval. There are subsequent fluctuations in the CAST coefficient that become increasing sharp after the 240-245 minute interval. This may be due to a transient period of stasis in differentiation shown in Figure 1.

## Data and Visualization Discussion

We have presented an analysis and visualization of cellular differentiation at a critical time period in *C. elegans* embryogenesis. The 200 to 400 minute interval is the time between the first appearance of terminally-differentiated cells outside of the germ line and the comma stage of development [20]. It is during the first part of this time period that the major differentiated cell categories are established. This has been done by looking at the ratio of developmental cells to terminally-differentiated cells, looking at the different cell families and the relative timing of their differentiation, and variation in timing within and between functional classes.

Looking between functional classes also reveals information about how larger-scale structures are built (e.g. nervous system). For example, Figure 6 shows the relationship between interneurons and neurons (A) and interneurons and hypodermal cells (B). In Figure 6A, the appearance of neurons is multimodal with respect to time (one early group and a larger latter group). By contrast, almost all interneurons appear after 275 minutes.

**FIGURE 6.**
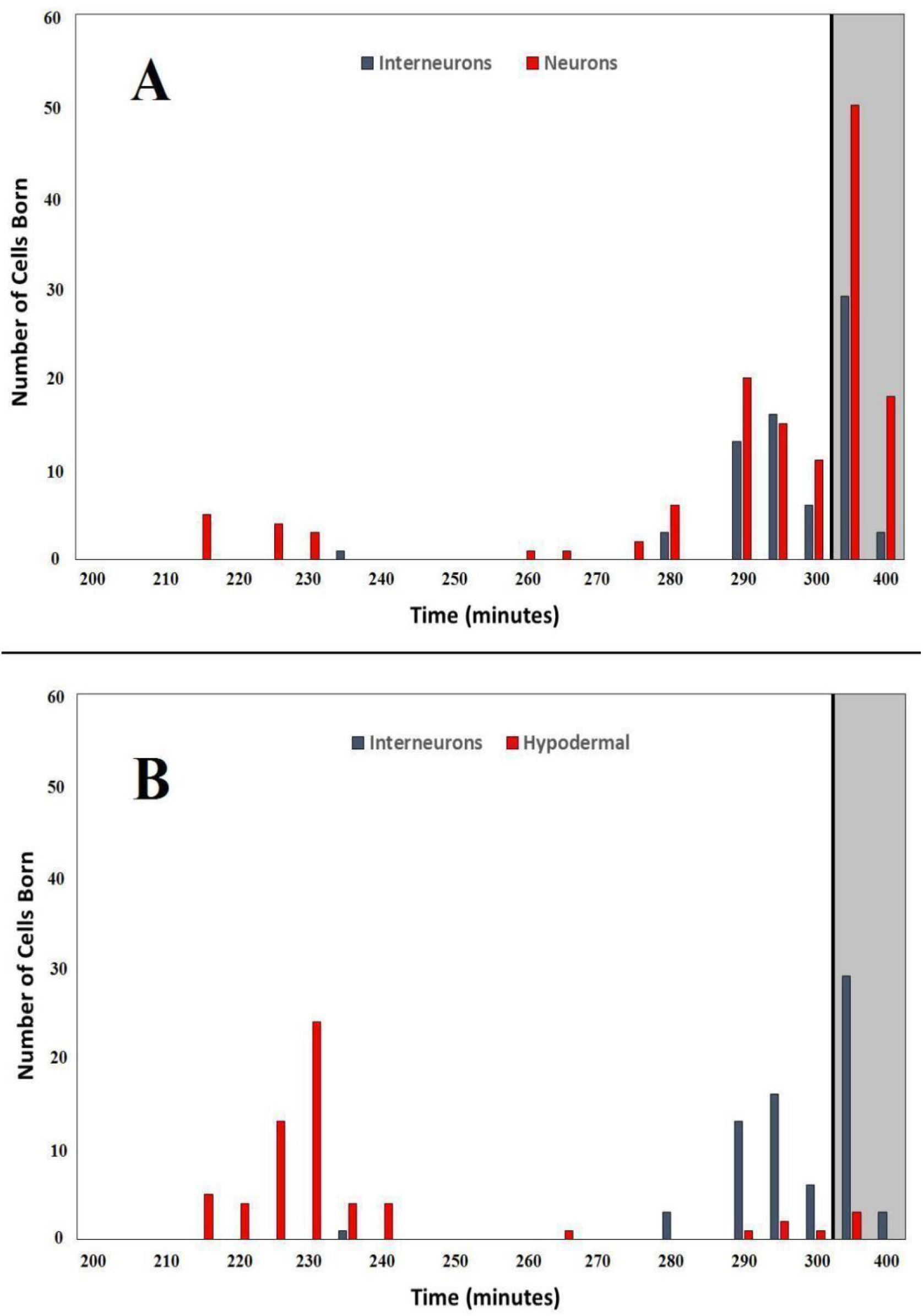
Histogram containing counts of types of cells born during a specific time interval (bins of size 10 except gray region denotes bins of size 50). Figure 6A: Interneurons (blue) vs. Neurons (red), Figure 6B: Interneurons (blue) vs. Hypodermal cells (red). Data collected from embryos raised at 25°C.

The timing of hypodermal cells is even more striking in comparison to interneurons as shown in Figure 6B. In this case, a large group of hypodermal cells appear before the sampled interneurons, while a smaller group of hypodermal cells appear alongside the sampled interneurons. These types of comparisons can provide clues as to the emergence of organs as well as other functional networks of cells (connectome).

The first consideration for further study is the behavioral relevance of structured differentiation. As autonomic (e.g. pharyngeal pumping) and other basic behaviors emerge from the developing embryo [21], we can ask questions regarding the minimal set of cells required for initiation of a given behavior, the appearance of cells essential to turning on that behaviour, and whether or not behavioral emergence involves more than terminally differentiated cells. Secondarily, the process of development can be represented as a spatiotemporal process (Figure 7). While this is foremost a data visualization problem, it is also critical in showing how the adult phenotype is modular with respect to developmental time. In a number of cases, we can observe a multitude of its components terminally differentiated well before the initiation of function.

**FIGURE 7.**
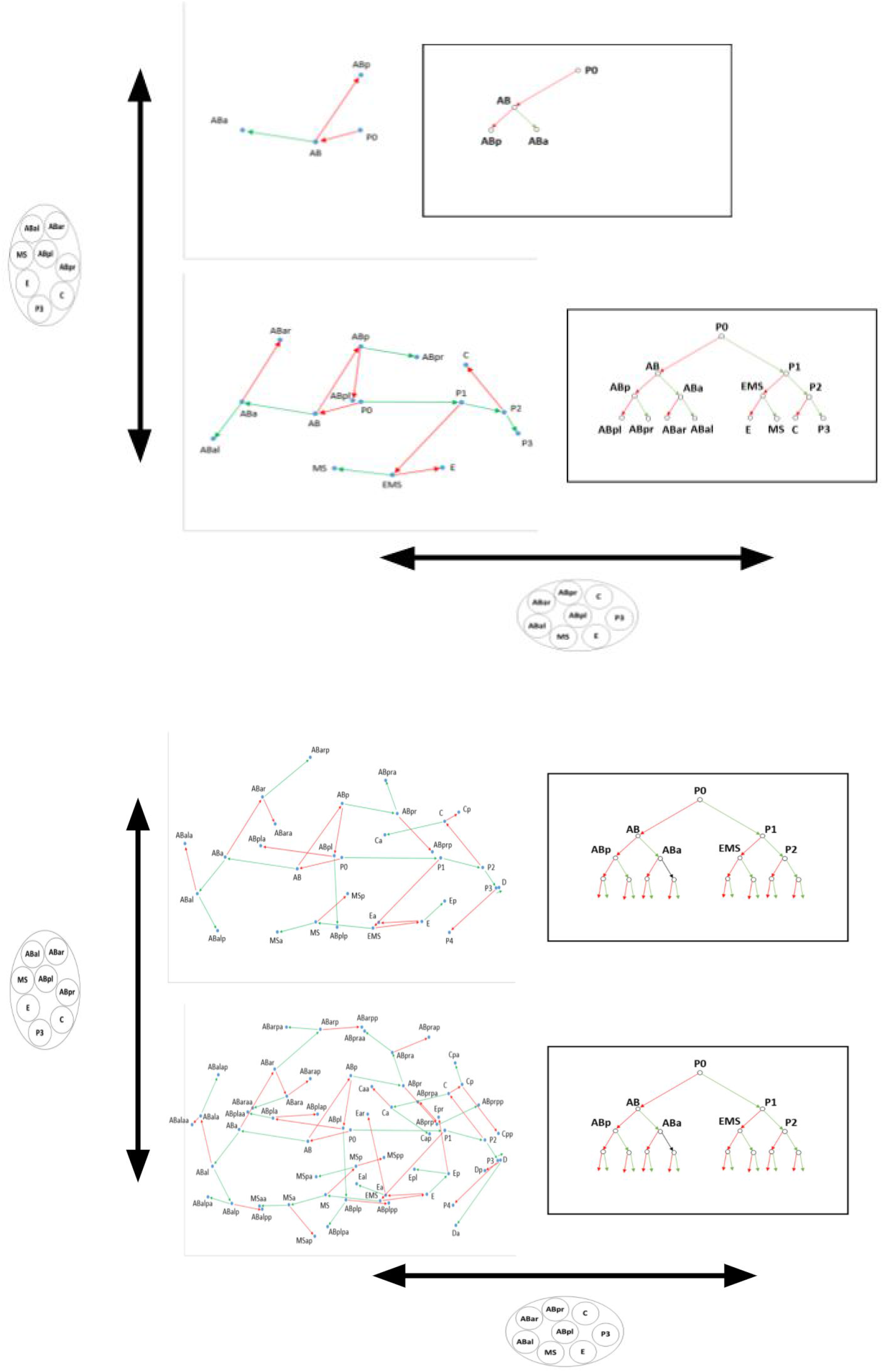
Differentiation maps for the *C. elegans* embryo at four different stages: 4-cell (upper top), 8-cell (lower top), 32-cell (upper bottom), 64-cell (lower bottom).

The visualization in Figure 7 is called a differentiation map which is based on the differentiation tree analysis of embryogenesis [12]. Each map is a 2-D representation of cell division as a spatial process. The extent of each differentiation map corresponds to the number of divisions in the lineage tree. Each cell is located by its position on the anterior-posterior (x) axis and the left-right (y) axis (in embryo units, AU). The lines between cells provide information about the change in position between a mother cell (e.g. AB) observed at time 0 and daughter cells (e.g. ABa, ABp) observed at time 1. Information about the GFP area tells us whether the line leading to either the smaller cell of the division (red) or larger cell of the division (green). Insets for each differentiation map shows the corresponding differentiation tree. For purposes of space, we truncated the 64-cell trees at 32 nodes (4 divisions).

Differentiation waves involve propagation of either a contraction or expansion of the apical surfaces of cells in a given epithelial tissue. In the case of mosaic development (such as in the case of *C. elegans*), tissues are replaced with individual cells [12]. In other words, an asymmetric cell division involves both a single-cell contraction wave, resulting in the smaller cell, accompanied by a single-cell expansion wave, resulting in the larger cell. An exception to this involves the small proportion of the cell divisions in *C. elegans* are symmetric, resulting in tissues containing two cells [12]. This set of rules allows us to bring regulative and mosaic development under one theory, the difference being that in regulative embryos tissues consist of many cells, whereas in mosaic embryos tissues consist of one cell.

### Synthesis of Data and Visualization

Structures describing the differentiation process (such as the differentiation wave) provide a means to determine the emergence of function in embryogenesis. In the model organism *C. elegans*, a deterministic developmental trajectory combined with available secondary data can be used to determine when terminally-differentiated cells appear and their relationship to both cell lineages and the adult phenotype [22, 23]. These data can provide insights into how movement and other behaviors first turn on, such as in cases where a specific cell is required for a generalized behavior or response [24]. In general, there is a great deal known about why the temporal emergence of *C. elegans* tissues and organs from terminally-differentiated cells is tightly regulated. However, a systems-level analysis and visualization of these cells could allow us to understand which cell types and anatomical features are necessary and/or sufficient for the emergence of autonomic behaviors and functional phenotypes.

In *C. elegans*, cell division patterns directly correspond to cell fate [25]. Furthermore, the timing and ordered emergence of cells making up a specific tissue or organ is highly regulated at the molecular level. Heterochronic timing and associated heterochronic genes are major drivers of *C. elegans* embryogenesis, particularly since the developmental process is more discrete than in vertebrates [26]. Cellular behaviors such as reorientation and contraction accompany the multi-step morphogenesis of various anatomical structures [27]. The coordination of cell division timing is a complex relationship related to developmental timing, and leads to asynchrony of divisions between sister cells [28]. The pace of cell division itself is an important regulator critical for the normal formation of tissues and organs [29]. The failure of normal development outside a specific temperature range, such as has been observed in amphibians [30], could be investigated in *C. elegans* at the single cell level.

This time-dependent type of single-cell developmental regulation has consequences for differentiated cells that comprise specific tissues and organs. For example, every cell has a unique pattern of transcriptional regulation in embryonic development [31]. The dynamic regulation of each developmental cell [32] leads to differentiated cells with diverse functions [31]. A key to better understanding the coordination of cellular differentiation in development is to look at differential transcription within and between cells [33]. The timing of cell division and differentiation events appear to influence which parts of a tissue or organ form before others and ensure proper function [34]. There is also a functional role for certain types of cells, which thus must be present at a certain stage of embryogenesis for proper anatomical function and the onset of behaviors. For example, glial cells are all purpose cells that play a critical role in the onset of movement and autonomic behaviors [35]. The presence, and more importantly absence, of actin molecules in cells that make up certain anatomical structures can affect their formation and function [36].

### Future Directions in Quantitative Analysis

This study serves as a first step towards constructing theories and mathematical applications based on quantitative measurements of developmental systems. Taking a relational view of embryonic development will allow for a common language to be used across species and different patterns of development. By understanding cell identities and the relations between them (division and differentiation), we can then proceed towards developmental composition. A model of compositional development allows us to combine quantitative measurement and analysis to produce an emergent, generative system. This in turn allows us to use more sophisticated theories and representational systems to describe developmental dynamics.

This quantitative approach that we rely upon is dependent upon technologies such as cell tracking and image segmentation. Future work will benefit from a number of advances in machine learning and computer learning techniques. For example, multiview methods [37] can be used to uncover the diverse characteristics of a microscopy time-series, and enhance further analysis using neural networks [38]. Using different viewpoints also allows us to use methods such as Topology Anchor Point Optimization to extract an estimation of 3-D scenes from 2-D images independent of initial image orientation or perspective [39]. Finally, we can use techniques such as group-based nuclear norm and learning graph models can improve image segmentation in ways that reveal even more information in the service of constructing theoretical models [40].

### Figures as Heuristics for Future Theory Construction

Before we discuss theory construction, we will interpret the main figures in this paper in terms of how they support broader insights into biological development. While these interpretations are heuristic in nature, they might serve to inform future theory construction and might be confirmable using computer simulations. In Figure 1, cellular divisions at this stage are synchronous, although proliferation rates are lower compared to first divisions in the embryo. The number of developmental cells is roughly the same, while the number of differentiated cells is increasing with each division of undifferentiated daughter cells. Figure 1 provides a visual hypothesis for why differentiated cell types start appearing at different times.

Figure 2 provides a time-series summary of terminal-differentiation events across multiple cell families. For this analysis, cells are grouped into functional families based on nomenclature. Families are also used in Supplemental Figure 2 (to analyze developmental cells) and Supplemental Figure 5 (to visualize the information content of terminally-differentiated cell families). These family groupings help us visually compare how cells playing a role in various functions are put into place over the course of developmental time, and how this enables functions to come on line. In the case of developmental cell families, we can examine their relative birth and differentiation as an early detection measure of large-scale differentiation events linked to the establishment of functional modules [41]. Comparison by family helps us to understand the emergence of cells supporting one function relative to cells supporting another function.

Figures 3, 4, and 5 are all heat maps that provide an alternative to the information shown in Figure 2. These heat maps (subsets of Supplemental Figure 3, which shows all cells in the embryo that differentiate before hatching) reveal the timing of terminal differentiation events on a single cell basis, and thus reveal heterogeneity across broader classes of functional annotation. In these figures, a broader functional classification is chosen over familial distinctions in order to make more direct pairwise comparisons of functional emergence. Figure 3 specifically looks at the birth of hypodermal versus interneuronal cells, while Figure 4 targets the relationship between newborn neuronal and synticial cells. Figure 5 is similar to Figure 4, looking at the emergence of muscle cells instead of neuronal cells and then comparing them to synticial cells. The latter cell type is particularly interesting as the development of synticium is key to developing tissues that are robust to elastic deformation and mechanical stress [42].

Figure 6 presents a time series summary of terminal differentiation expressed as a histogram. Each part of Figure 6 features a pairwise comparison between three cells types: Figure 6A (a comparison of neurons and interneurons) is relevant to the assembly and emergence of the connectome [43, 44], while Figure 6B (a comparison of hypodermal and interneuronal cells) is a restatement of Figure 3. Comparing neurons and interneurons demonstrates that these different types of neuronal cells are born at similar times in development, which may be both a functional necessity and a product of developmental constraints (descending from similar developmental sublineages).

Figure 7 demonstrates a new means of data representation: the differentiation map. Differentiation maps are a bivariate graph of spatial trajectories, which represent angles of migration upon cell division inferred from discrete measurements of mother and daughter cell centroids. Maps generated from early *C. elegans* embryogenesis demonstrate great spatial variability across the anterior-posterior and left-right axes of the embryo. There exist regions of density as well as spatial locations where cells tend not to move. This may have relevance to understanding the behavior of cells in development, along with early physical constraints expressed in different sublineages. Information expressed through differentiation maps can also be applied to lineage tree analysis, and understanding complex network topologies.

In the next section, we will consider theory construction as a means to guide research in a manner different from standard hypothetico-deductive investigation. In many cases, and in high-throughput computational biology in particular [45], the formulation of hypotheses can work counter to discovery [46]. Yet exploratory quantitative endeavors are not particularly effective at conveying understanding. Most computational “insight” is largely utilitarian in nature. Considering the process of theory construction can help bridge this gap. Drawing from the allied activity of model building, it is clear that two factors are particularly crucial to the interpretation of data. The first factor involves deciding which components of the system under investigation should be represented in your model [47]. As theories can be both phenomenological and predictive in nature, choosing some components over others is often essential. Secondly, properly representing the order of operations is important for refining a story of the data [48]. Rather than propose a grand theory functional emergence in *C. elegans* development based on these data, in the next section we will consider how the philosophy of theory construction can more effectively help us interpret quantitative data analyses such as those presented in this paper.

### Revisiting Theory Construction

Physics is often viewed as the gold standard of successful theory construction, although this is a misconception for reasons we will not emphasize here. Biology presents a problem distinct from physics in that theory is often not explicitly predictive. While this can be used as a rationale to not engage in the development of theory, attempts at theory construction can nevertheless be highly useful. The distinction between theories of biology and physics involve historical chance, heterogeneous environmental effects, and non-linear behaviors resulting from adaptive processes [48]. Thus, for our purposes, model building from data can be an integral part of theory construction.

Eisenhardt and Graebner [49] propose using a case-based approach to create theoretical constructs or propositions. Case studies do not need to be quantitative, but must provide rich descriptions of particular instances of a phenomenon. In the developmental biology context, cases can be based on a variety of data sources or detailed computational representations. Based on knowledge acquisition from case studies, we can also build a cycle of description leading from initial observation to testing in the empirical world [50]. According to this view, initial observations lead to descriptive models, which are then fleshed out into explanatory frameworks. These frameworks are subsequently tested (or deployed) in the experimental realm. In the realm of Computational Developmental Biology [51], representational frameworks and simulation often takes the place of testing/deployment.

Theory construction can resemble an incremental process, but is often a chaotic process driven by shifts in conceptual understanding and changes in the epistemic basis of a given scientific field [52, 53]. Such shifts are often the product of reaching certain thresholds of knowledge and conceptual understanding [54], which once reached results in an avalanche of theoretical development. This is broadly characterized by so-called paradigm shifts [55], but is often driven by changes in methodology, particularly technological advances in measurement and computing. In contexts where we have sparse quantitative data regarding relational causal mechanisms, philosophy can assist in the prospect of building robust theories. An example of this is stemness [56], where data regarding the classification of cell types [57] can be quite difficult to interpret [58]. In this case, methods such as conceptual analysis and argumentation allow us to propose knowledge and conceptual frameworks in support of predictive theories [59].

### Towards a category theory of development

As demonstrated by our data analysis, the process of cellular differentiation leads to two distinct categories: developmental cells that are not differentiated, and terminally-differentiated cells that exhibit some (but not all) attributes of cells found in the adult. But we can also have duality of categories in terms of axial differentiation (e.g. anterior-posterior), or in terms of emerging asymmetry (left-right), the interpretation of which is aided through the application of differentiation maps (Figure 7). More broadly, the data analyses provided here reveal the basic structure necessary for a theory of functional emergence in development.

One candidate framework for theory construction is applying category theory [60] to the problem of functional emergent in development. A theoretical representation of developmental plasticity, even in a so-called deterministic organism such as *C. elegans*, requires flexible categories amenable to mathematical operators as well as being robust to dynamical systems [61]. A category theory interpretation requires us to define *objects* and *functions* as the basis for functional categories and relationships between categories, respectively. Examples of objects include terminally-differentiated cell families and birth-time cohorts. Mathematical functions that constitute formal biological categories include cell division events, changes in spatial location, and transitions of an object through the developmental process. The categories as defined by category theory do not exist in isolation, and in fact can intersect in a number of ways.

An additional advantage to thinking about the morphogenetic process in a category theoretic fashion is the ability to utilize a concept called compositionality [62]. In this case, compositionality allows us to piece together events from trends in familial classifications. This can range from the number of cells differentiated at a given point in time, or proportions of cells in a given functional class. Extending the diagrammatic relationships shown in Figure 7, we can propose that there is the value in using concepts such as natural transformations [63] as a way to capture categorical transformations due to cell division and differentiation. Operating at a higher level of complexity, natural transformations provide the ability to reconfigure categories of functors while preserving their structure. This is useful in the developmental application domain, where we might view the role of cell differentiation as a mathematical function between the categories of developmental and terminally-differentiated cells, and the role of natural transformations as a means to represent variation in the differentiation process across various functional cell types.

## Acknowledgments

We would like to acknowledge feedback from the OpenWorm and DevoWorm communities, particularly Drs. Stephen Larson, George Mikhailovsky, and senior contributors at the OpenWorm Foundation.

## SUPPLEMENTAL FIGURES

Supplemental Figure 1. A comparison of developmental and terminally-differentiated cell counts for 50-minutes intervals. Data collected from embryos raised at 25 °C.

Supplemental Figure 2. The number of developmental cells alive for each sublineage (AB, MS, C, D, and E) at 20 minute time intervals for 200-400 minutes of embryogenesis. Panel A shows AB and MS, while panel B shows C, D, and E. Data collected from embryos raised at 25 °C.

Supplemental Figure 3. Heat map showing the emergence of terminally differentiated cells in *C. elegans* from 200 to 400 minutes of embryogenesis. Emergence of Terminally Differentiated Cells in each cell type class, relative to the 400 minutes of embryogenesis. The 200 to 300 minute period is sampled at 5-minute intervals, while the 300 to 400 minute period is sampled at 30-minute intervals. Bottom of the figure is labeled with the corresponding developmental stages and images of the embryo during select times. Data collected from embryos raised at 25 °C.

Supplemental Figure 4. Histogram containing counts of types of cells born during a specific time interval (bins of size 10 except where noted). Figure 5A: Interneurons (blue) vs. Neurons (red), Figure 5B: Interneurons (blue) vs. Hypodermal cells (red). Gray region denotes bins of size 50. Data collected from embryos raised at 25 °C.

Supplemental Figure 5. Information content for each terminally-differentiated cell family, based on a hierarchical clustering analysis. Information content (blue bars, left axis) is compared to the number of cells in each family (red bars, right axis). Data collected from embryos raised at 25 °C.

Supplemental Figure 6. A time-series of CAST coefficients for 200 to 400 minutes of *C. elegans* embryogenesis. Time intervals from 200 to 300 minutes are five minutes in length; time intervals from 300 to 400 minutes are fifty minutes in length (denoted within gray region). Data collected from embryos raised at 25 °C.

## SUPPLEMENTAL FILES

Supplemental File 1. Terminally-differentiated cell nomenclature identities and annotations by developmental birth time (min).

Supplemental File 2. Table of number of cells born at a specific developmental birth time sampling point (min) for five distinct somatic cell types.

Supplemental File 3. Table of somatic cell types by family, class, and developmental birth time (min).

Supplemental File 4. Table of cell families by number of family members and average developmental birth time (min).

Supplemental File 5. Pairwise alignments (per pairs of birth time sampling points) and calculation of alignment scores for CAST analysis.

